# Fine-mapping and QTL tissue-sharing information improve causal gene identification and transcriptome prediction performance

**DOI:** 10.1101/2020.03.19.997213

**Authors:** Alvaro N. Barbeira, Yanyu Liang, Rodrigo Bonazzola, Gao Wang, Heather E. Wheeler, Owen J. Melia, François Aguet, GTEx Consortiumt, Kristin G Ardlie, Xiaoquan Wen, Hae K. Im

## Abstract

The integration of transcriptomic studies and GWAS (genome-wide association studies) via imputed expression has seen extensive application in recent years, enabling the functional characterization and causal gene prioritization of GWAS loci. However, the techniques for imputing transcriptomic traits from DNA variation remain underdeveloped. Furthermore, associations found when linking eQTL studies to complex traits through methods like PrediXcan can lead to false positives due to linkage disequilibrium between distinct causal variants. Therefore, the best prediction performance models may not necessarily lead to more reliable causal gene discovery. With the goal of improving discoveries without increasing false positives, we develop and compare multiple transcriptomic imputation approaches using the most recent GTEx release of expression and splicing data on 17,382 RNA-sequencing samples from 948 post-mortem donors in 54 tissues. We find that informing prediction models with posterior causal probability from fine-mapping (*dap-g*) and borrowing information across tissues (*mashr*) lead to better performance in terms of number and proportion of significant associations that are colocalized and the proportion of silver standard genes identified as indicated by precision-recall and ROC (Receiver Operating Characteristic) curves. All prediction models are made publicly available at predictdb.org.

**Author summary:** Integrating molecular biology information with genome-wide association studies (GWAS) sheds light on the mechanisms tying genetic variation to complex traits. However, associations found when linking eQTL studies to complex traits through methods like PrediXcan can lead to false positives due to linkage disequilibrium of distinct causal variants. By integrating fine-mapping information into the models, and leveraging the widespread tissue-sharing of eQTLs, we improve the proportion of likely causal genes among significant gene-trait associations, as well as the prediction of “ground truth” genes.

## Introduction

Transcriptome studies with whole genome interrogation characterize genetic effects on gene expression traits. These mechanisms help elucidate the function of loci identified in genome-wide association studies (GWAS) by identifying potential causal genes that link genetic variation with complex traits [1–5] {Albert:2015fx, Aguet2019, Huckins:2019ix, Mancuso:2018fv, Gusev:2018dy}.

In particular, the Genotype-Tissue Expression (GTEx) project [2] {Aguet2019} has sequenced whole genomes from 948 organ donors and generated RNA-seq data across 52 tissues and 2 cell lines. Results and tools derived from this comprehensive catalog of transcriptome variation have enabled a myriad of applications such as drug repurposing 6 {So2017} and clinical discoveries in cancer susceptibility genes [7] {Wu2018}, to name a few.

The general consensus that many noncoding variants associated with complex traits exercise their action via gene expression regulation has motivated the development of imputed transcriptome association approaches such as PrediXcan [8, 9] {Gamazon2015, Barbeira2018}, TWAS/FUSION [10] {Gusev:2016ey} and UTMOST [11] {Hu2019}. In essence, these methods predict gene expression traits based on individuals’ genotypes and test how these predictions correlate with complex traits.

Reliable prediction models for gene expression traits are key components of imputed transcriptome association studies. Given the predominantly sparse genetic architecture of gene expression traits [12] {Wheeler2016} and overall robustness and performance [3, 13] {Huckins:2019ix, Fryett:2018bg}, Elastic Net [14] {Friedman2010GLMNET} has become the algorithm of choice for predicting transcriptome variation.

Despite Elastic Net’s many advantages such as robustness and sparcity, we hypothesized that transcriptome imputation can be improved by leveraging biologically-informed methods. Recent efforts [11] {Hu2019} have exploited the high degree of eQTL sharing across tissues [15] {GTEx2017} by leveraging cross-tissue patterns in the broad GTEx panel to improve prediction performance, more notably in tissues with small sample sizes. Also, important methodological progress in fine-mapping [16, 17] {Wen2017, Wang2018} and an adaptive shrinkage method that improves effect size estimates across multiple experiments [18] {Urbut2019} provide opportunities to further improve quality of downstream associations.

In this article, we analyze different transcriptome prediction strategies and compare their strengths both in prediction performance and downstream phenotypic associations. Proximity and linkage disequilibrium (LD) between distinct causal variants can lead to non causal associations between predicted expression and complex traits [9, 19] {Barbeira2018, Wainberg:2019kq}. Since the ultimate goal of imputed transcriptome studies is to identify causal genes, our main focus here is to improve discoveries with less emphasis on expression prediction performance. We also applied the same model building techniques to alternative splicing traits quantified with Leafcutter [20] {Li:2018cy}. We make all results, prediction models and software available to the research community.

## Results

To identify optimal techniques for transcriptomic imputation, we have built models to predict genetically regulated expression (GREx) using four different approaches on GTEx expression and splicing data (release version 8). To reduce LD misspecification problems, most apparent when applying summary statistics-based versions of PrediXcan on GWAS of European populations, we used only European samples.

We restricted the analysis to genes that are annotated as protein coding, lncRNA, and pseudogenes in GENCODE version 26 [21] {Frankish2019}. We included 49 different tissues with sample sizes ranging from 65 (Kidney Cortex) to 602 (Muscle Skeletal).

The first strategy used the Elastic Net [14] {Friedman2010GLMNET} algorithm to compute predictions as described previously in [8, 12] {Gamazon2015, Wheeler2016}. For every gene available in each tissue, this strategy used variants from the HapMap CEU track in a window ranging from 1Mb upstream of the transcription start site to 1MB downstream of the transcription end site as explanatory variables. Only those models achieving thresholds of cross-validated correlation *ρ* > 0.1 and prediction performance p-value < 0.05 were kept. We will refer to this family as the **EN-M** models.

The second strategy used CTIMP (Cross Tissue gene expression IMPutation) [11] {Hu201} CTIMP uses a regularized, generalized linear regression algorithm to fit expression from different tissues simultaneously. CTIMP optimizes a cost function including a within-tissue Lasso penalty and a cross-tissue group Lasso penalty, thus inheriting Lasso-like behaviour that is less sparse than Elastic Net. We used the same variants from the EN-M strategy (HapMap CEU track, same windows around each gene), and identical correlation threshold (*ρ* > 0.1) and cross-validated prediction performance threshold (*p* < 0.05) to accept models. We will refer to this family as the **CTIMP-M** family. We verified that this method’s performance is not significantly improved by using all available GTEx variants, as explained in the supplementary material.

The third strategy used the posterior inclusion probability (PIP) of a variant being causal for gene expression as estimated by the Bayesian fine mapping method *dap-g* (Deterministic Approximation of Posteriors) [22] {Wen2016}. First, for every gene, we restricted to variants with posterior inclusion probabilities PIP > 0.01. Since *dap-g* clusters variants by their LD, we kept the variant with highest PIP from each cluster to avoid redundant explanatory variables. Then, the selected variants were fed into the Elastic Net algorithm, scaling each variant’s effect size penalty by a factor of 1 − PIP (i.e. more likely variants are less penalized). Only those models achieving good enough cross-validated prediction performance (p-value< 0.05) and correlation (*ρ* > 0.1) were kept. We will refer to this family as **DAPGW-M** (*dap-g* weighted). As discussed later, the cross-validated prediction performance of this approach can’t be fairly compared to EN-M and CTIMP-M because the pre-selection of fine-mapped variants is based on the same underlying data.

The fourth strategy used *mashr* (Multivariate Adaptive Shrinkage in R) [18] {Urbut2019} effect sizes from variants selected by *dap-g* as in the **DAPGW-M** approach. More specifically, fine-mapped variants were selected as in the **DAPGW-M** approach but the weights were obtained by applying *mashr* to the marginal effect sizes and standard errors from the GTEx eQTL analysis [2] {Aguet2019}. Unlike the previous methods, this approach does not fit into a cross-validation strategy and therefore lacks a natural prediction performance measure. Only eGenes with at least one cluster of variants achieving *dap-g* PIP> 0.1 were kept. We will refer to this family as **MASHR-M**.

We did not consider the BSLMM family of methods for transcriptome prediction. These models contain both a sparse and a polygenic component. The latter is likely to induce LD contamination [9] {Barbeira2018} and doesn’t reflect the sparse architecture of expression traits [12] {Wheeler2016}.

We also applied the EN-M and MASHR-M methods to alternative splicing quantification from LeafCutter [20]{Li:2018cy} and made them readily available to the research community. These models were extensively used in [2]{Aguet2019} and [23]{GTEx-GWAS-Companion}.

### Summary of models

Given the differences in computational approach, not all prediction strategies generated models for every available gene-tissue pair. As can be seen in Fig. 1-A, EN-M yielded the smallest number of valid models, for 281,848 gene-tissue pairs. CTIMP-M produced 340,104 valid models, 21% more than EN-M, as expected from its integration of multiple tissues’ information.

**Fig 1.**
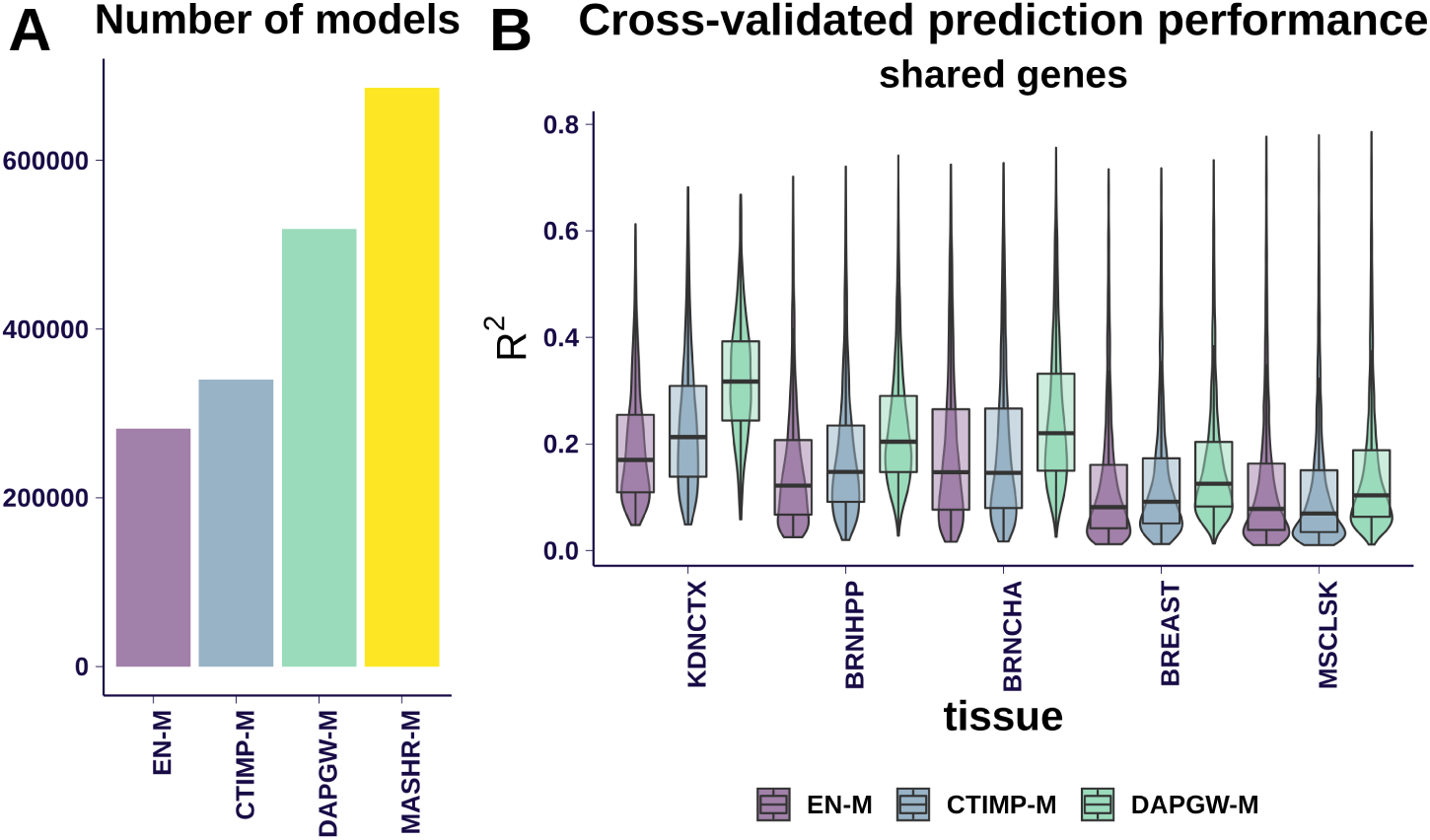
Models summary. **Panel A** shows the number of models generated for protein coding genes, pseudo genes and lncRNA across the four strategies. MASHR-M displayed the largest number of generated models. **Panel B** compares prediction performances for gene-tissue pairs present in all four strategies, at 5 different tissues ordered by sample size. CTIMP-M performed better than EN-M in tissues with smaller sample size. DAPGW-M is presented for illustration purposes; since it included an additional variable selection step using the same underlying data, it cannot be fairly compared to EN-M and CTIMP-M. MASHR-M doesn’t have a prediction performance measure. The intersection of gene-tissue pairs across the 4 strategies is mostly defined by Elastic Net, the smallest set. 82% of Elastic Net models make up the intersection available to all strategies. **Tissue abbreviations and sample size:** KDNCTX: Kidney - Cortex, n=65; BRNHPP: Brain - Hippocampus, n=150; BRNCHA: Brain - Cerebellum, n=188; BREAST: Breast - Mammary Tissue, n=337; MSCLSK: Muscle - Skeletal, n=602

Fine-mapping-based methods generated even more models: 518,537 from DAPGW-M (84% more than EN-M) and 686,241 from MASHR-M (143% more than EN-M). Please note that given the different criteria used to accept a model as valid, simple counts of available models should not be considered a measure of performance.

We show the distribution of cross-validated prediction performances in Fig. 1-B. We include 5 representative tissues ordered by increasing sample size (kidney, brain - hippocampus, brain - cerebellum, breast, skeletal muscle). In order to perform a uniform comparison, we used only gene-tissue pairs available to all model families. CTIMP-M showed better prediction performance than EN-M on tissues with smaller sample size, but performed similarly on tissues with larger sample sizes. We attribute this to CTIMP’s design, which leveraged all existing samples’ genotypes in the tissues of smaller expression sample size. MASHR-M models had no natural prediction performance measure and thus are excluded from these panels. DAPGW-M is presented for completeness but its comparison to EN-M and CTIMP-M is unfair. We show in Sup. Fig 1 the cross-validated prediction performances for all genes in each family.

### Finemapping improves expression prediction in independent datase

Next, we sought to validate the models’ predictions in an independent RNA-seq dataset. We analyzed data from the the GEUVADIS project [24] {Lappalainen2013}, which includes 341 samples of European ancestry with genotype and LCL (lymphoblastoid cell lines) expression data. We predicted expression using GTEx LCL models from the 4 strategies, and compared with measured expression levels. Fig. 2-A shows the number of genes that each family was able to predict. DAPGW-M and MASHR-M had the largest number of predictable genes, followed by CTIMP-M and EN-M.

**Fig 2.**
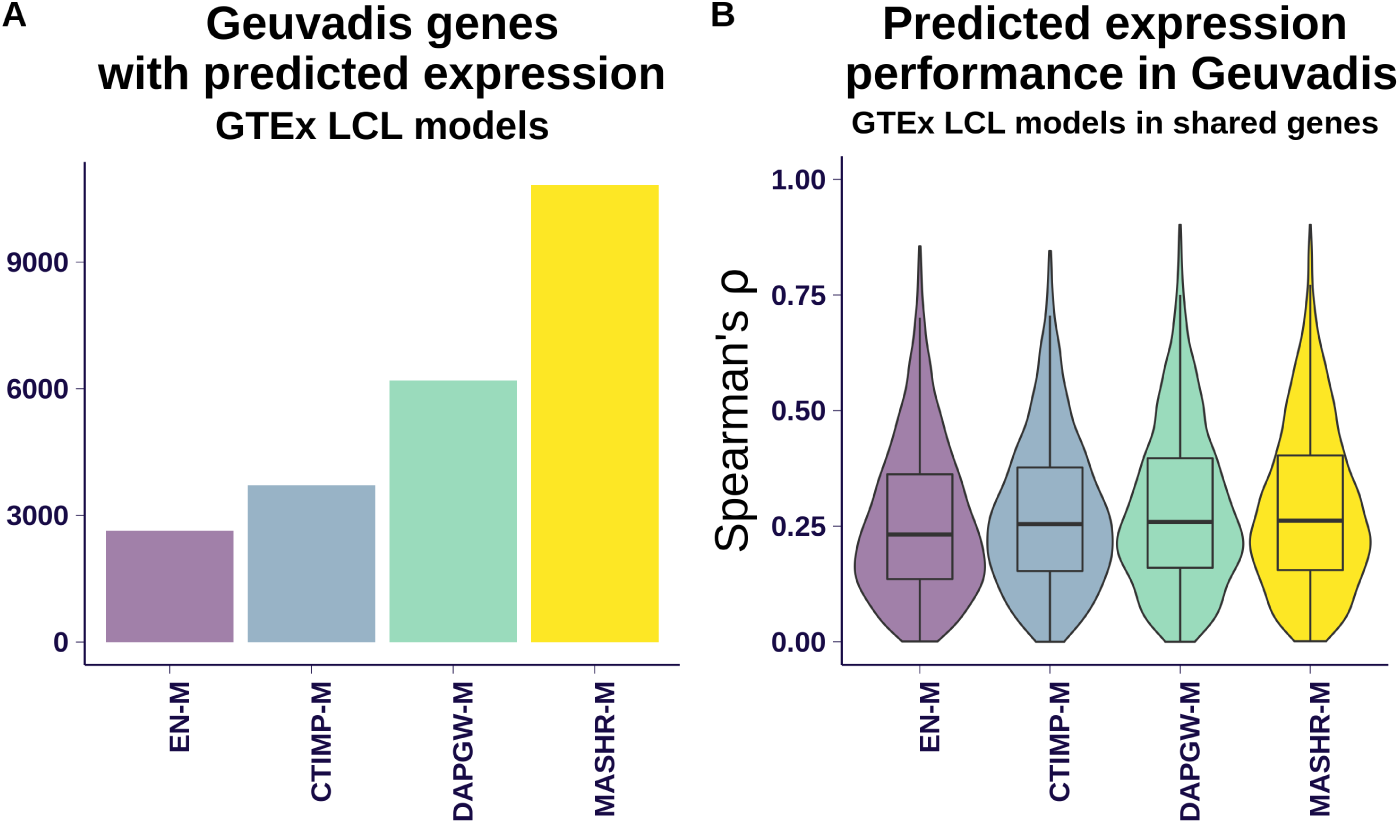
Validation in a separate expression cohort. Panel **A** shows the number of genes predicted in GEUVADIS cohort using the LCL models from each of the four strategies. MASHR-M had the most models available, followed in decreasing order by DAPGW-M, CTIMP-M and EN-M. Panel **B** shows the distribution of prediction performances (Spearman *ρ*)for genes available to all four families. DAPGW-M and MASHR-M performed slightly but consistently better than EN-M and CTIMP-M. We attributed the small differences to the GTEX LCL tissue having a small sample size (n=115 individuals), much lower than the 341 available in GEUVADIS. Also, the intersection of genes available to all 4 strategies is dominated by those present in Elastic Net, the smallest set; and genes that can be modelled with Elastic Net tend to be the ones with less complicated patterns of variation.

To compare prediction performances, we used Spearman’s rank correlation coefficient *ρ* as a robust measure that handles the scale and complexity differences between real GEUVADIS expression data and predicted expression levels. Fig. 2-B shows the distribution of prediction performance (Spearman’s *ρ*) for genes present in all four methods on the LCL tissue. We observed that all four families achieved similar levels of performance, with DAPGW-M and MASHR-M faring slightly but consistently better than EN-M and CTIMP-M.

We attribute the smaller performance differences to low power, since GTEx LCL tissue has a sample size of n= 115 individuals, much lower than the 341 available in GEUVADIS.

### Fine-mapping improves number and colocalization of associations

Next, we assessed whether any of these models perform better at identifying causal genes. We considered the number and proportion of colocalized genes among the significant ones as measures of association quality.

We used the four families of models to correlate predicted expression with 87 phenotypes through 49 tissues using the summary version of PrediXcan. Results of applying the EN-M models to GWAS summary statistics, harmonized and imputed to GRCh38 [25] {Schneider2017hg}, were presented in [2] {Aguet2019}. In this section, we say that a gene-tissue pair is significant if it achieves a p-value below the Bonferroni-corrected threshold (0.05*/*number of gene-tissue pairs) within each trait.

We used *enloc [16]* {Wen2017} results published in [2] {Aguet2019} to assess the colocalization status of GWAS and transcriptomic traits as evidence for a shared underlying mechanism. Briefly, *enloc* computes the “regional colocalization probability” (rcp) that a trait shares causal variants with a gene’s expression (or an intron’s splicing quantification), within a GWAS region and the overlapping gene’s cis-window. We say that a gene-tissue pair is “colocalized” with a trait if it achieves an *enloc* regional colocalization probability *rcp* > 0.5. Note that *rcp* <= 0.5 should not be interpreted as a false association; rather, it only means that there is not enough evidence of colocalization. See discussion on the conservative nature of colocalization approaches in [23] {GTEx-GWAS-Companion}.

We say that a gene-tissue pair that is both significant and colocalized is a “prioritized” detection or candidate. To simplify interpretation of results across multiple tissues, we count the number of unique genes among the prioritized gene-tissue pairs for each trait.

We found that MASHR-M tipically yields more candidate genes. We display the numbers of detections for each trait in Fig. 3, through Q-Q plots comparing MASHR-M to the other the model families. We observe in Fig. 3-A that the fine-mapping informed families of models, DAPGW-M and MASHR-M, yielded a similar number of candidates per trait, consistently larger than EN-M and CTIMP-M. When comparing the fraction of colocalized genes among significant genes (3-B), MASHR-M performs better than the other 3 families. In general, we observed that associations obtained through both DAPGW-M and MASHR-M models tend to agree (see Supplementary Figure 2 as an example).

**Fig 3.**
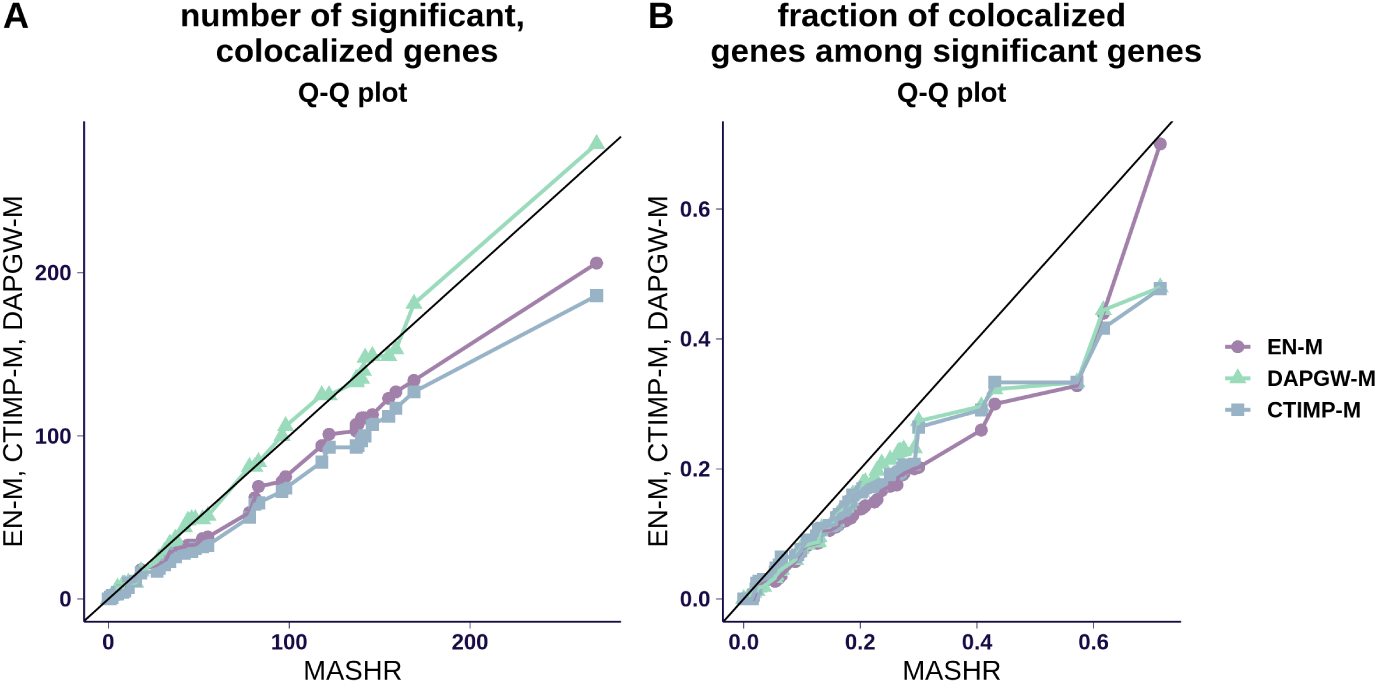
PrediXcan associations across 87 traits. **Panel A** shows a Q-Q plot for the number of colocalized, significant genes per trait. Fine-mapping-informed models (DAPGW-M and MASHR-M) achieved similar numbers of colocalized detections, both slightly higher than EN-M and CTIMP-M. **Panel B** shows a Q-Q plot for the fraction of colocalized genes among significant genes per trait. MASHR-M’s distribution is shifted towards higher proportions than the other families. We say a gene is significant if it achieves a Bonferroni-adjusted threshold of 0.05*/*number of available gene-tissue pairs, in at least one tissue. Likewise, we say a gene is colocalized if it achieves *enloc rcp* > 0.5 in any tissue. We say a gene is a candidate or “prioritized” detection if it is both significant and colocalized in any tissue.

We were thus led to favor MASHR-M, which produced the largest number of models, with superior number of colocalized, significant associations as well as higher proportions of colocalized associations among significant genes.

*Enloc* relies on the *dap-g* algorithm itself as a component, so that the fraction of colocalized genes could have been biased towards *dap-g* informed methods. To make sure that the use of *dap-g* is not driving the improved colocalization rate of MASHR-M over th other strategies, we verified the performance using another colocalization method, *coloc [26]* {Giambartolomei2014}.

We observed that MASHR-M still had a better rate of colocalization among significant associations, albeit with smaller differences as can be seen in Supplementary Figure 3. This is probably in part due to *coloc*’s reduced power and limiting assumption of a single causal variant (see [23]{GTEx-GWAS-Companion} for details).

### Finemapping improves identification of silver standard genes

As an independent way to assess each prediction strategy’s ability to identify causal genes, we framed the problem as one of causal gene prediction and use standard prediction performance measures such as Receiver Operating Charasteristc (ROC) and Precision-Recall (PR). This avoids using an ad-hoc significance or colocalization thresholds.

As proxies for causal genes, we leveraged two different “silver standards” as described in Barbeira et al. [23] {GTEx-GWAS-Companion}. The first one, based on the OMIM (Online Mendelian Inheritance in Man) database [27] {amberger:2019}, features 1592 known gene-trait associations. The second one is based on rare variant association studies [28–30] {marouli:2017, liu:2017, locke:2019} and contains 101 gene-trait associations.

We restricted our analysis to gene-trait pairs in the vicinity of the corresponding traits’ GWAS loci since we did not expect any of the methods to detect reliable signals elsewhere. We used approximately-independent LD regions [31]{berisa:2016} to define vicinity.

Using absolute values of z-scores as association score for each strategy, we assessed their ability to ‘predict’ the silver standard gene-trait associations. We show in Fig. 4 the ROC and Precision-Recall curves on OMIM- and rare variant-based silver standards.

Using the OMIM-based silver standard (Fig. 4-A and -C), we observed that MASHR-M strategy outperforms the other strategies, with DAPGW-M a close second. Thus we concluded that MASHR-M models are better equipped for detecting known genes in the extreme regulatory case of Mendelian diseases, reinforcing our choice of MASHR-M as the best option.

**Fig 4.**
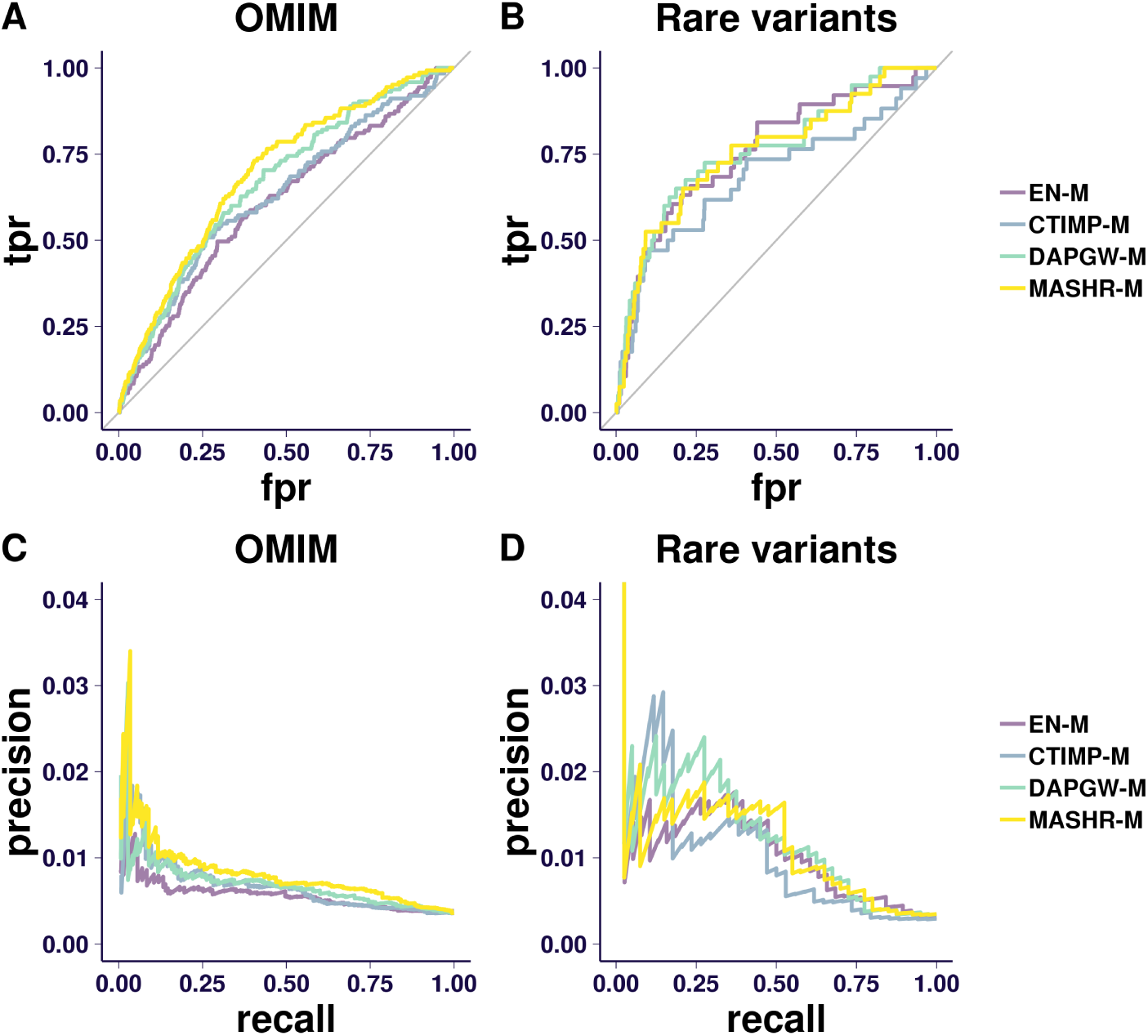
ROC and PR curves. **Panel A** shows the Receiver Operating Characteristc (ROC, plotting true-positive ratio to false-positive ratio) curve for the OMIM silver standard. We observe that MASHR-M models outperforms the other strategies, with DAPGW-M a close second. **Panel B** shows the ROC curve for the rare-variant-based silver standard. We observe that all strategies perform better than taking a random choice. However, this silver standard is too limited to properly distinguish between strategies. **Panel C** shows the Precision-Recall (PR) curve for the OMIM silver standard. MASHR-M performs better than the other strategies in general, but precision becomes a noisy measure towards lower recall ranges. **Panel D** shows the PR curve for the rare-variant-based silver standard. The precision measure is too unstable to draw any conclusions. The OMIM silver standard not only validates the 4 proposed model strategies as a consistent approach to detect causal genes, but provides additional evidence of MASHR-M’s superiority. The second silver standard, based on rare variants, is too limited to conclude anything beyond a high-level validation of all 4 families.

Using the rare-variant-based silver standard (Fig. 4-B and -D), we observed that all four strategies are able to detect known causal genes. However, the limited size of this standard did not allow us to distinguish between the four families.

### Importance of harmonization and imputation of missing summary statistics

The prediction models’ usefulness depends on the availability of their variants in the GWAS of interest. Publicly available GWAS use different sequencing and genotyping techniques, based on different genotype imputation panels and human genome release versions, so that the lists of available variants vary wildly across traits. Thus, a GWAS might lack particular variants from a prediction model, so that the model can’t properly infer variation patterns as shown in [32] {Barbeira2019}. Since many fine-mapped variants in the GRCh38-based GTEx study can be absent in a typical GWAS, we sought to assess the impact of variant compatibility in real applications.

We compared S-PrediXcan results from MASHR-M models on 69 publicly available GWAS with two preprocessing schemes:

1. Harmonization of variants by mapping genomic coordinates between human genome assemblies, and filtering for matching alleles (“Harmonization” for short)
2. Imputation of missing summary statistics (“Imputation” for short) on harmonized GWAS.

The 69 traits included in this analysis are those among the 87 traits not belonging to the Rapid GWAS project, to prevent the highly homogeneous Rapid GWAS datasets from dominating comparisons.

We show in Fig. 5 the effect of these preprocessing schemes on various performance metrics, segregated by human genome release version (hg17, hg18, hg19).

**Fig 5.**
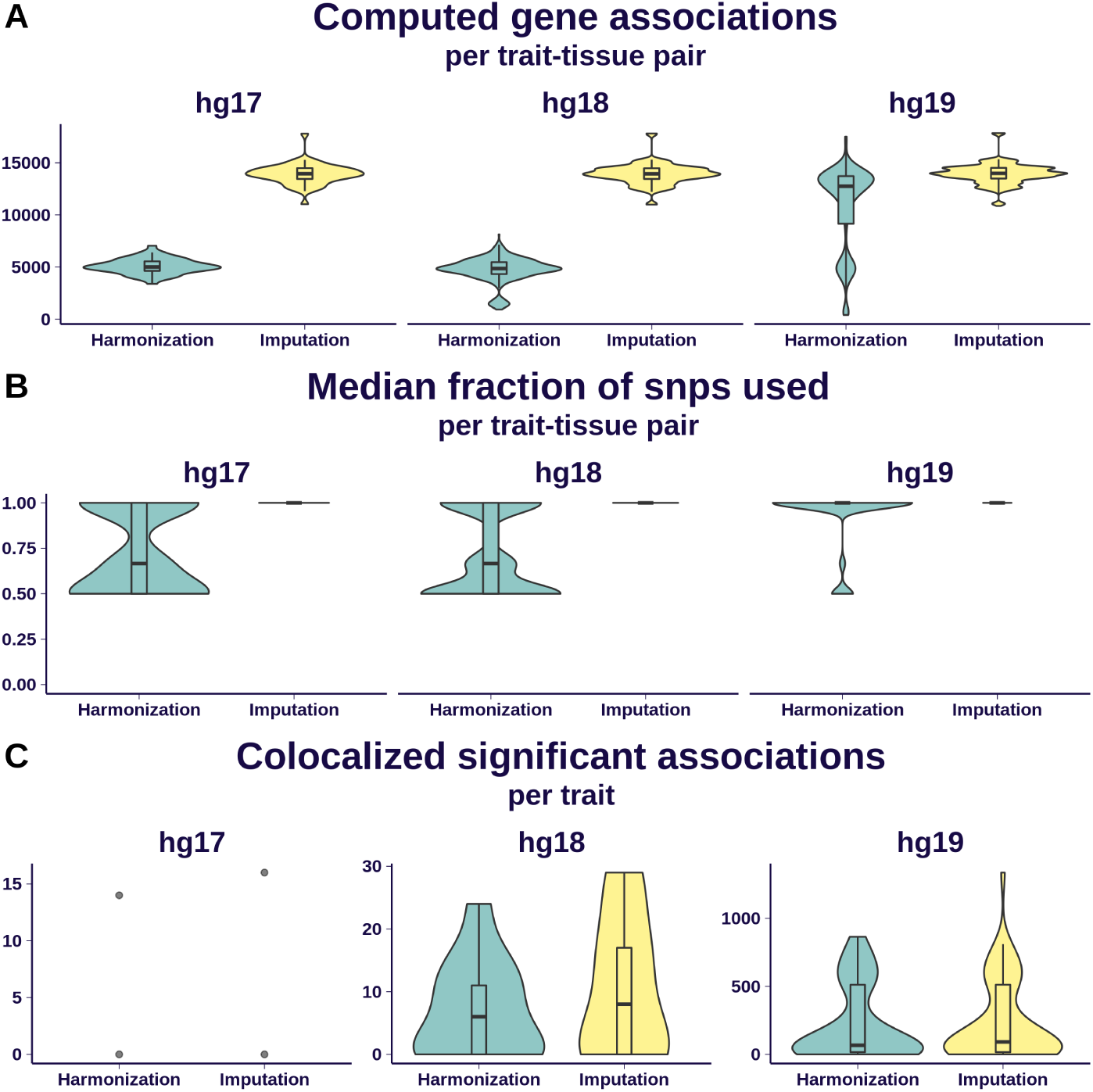
Effect of imputation on association quality. We display here a comparison of S-PrediXcan results from MASHR-M models on 69 GWAS traits using two different pre-processing schemes: simple harmonization of GWAS variants to GTEx’s, and additional imputation of missing summary statistics. Results are grouped by the different human genome release versions underlying each GWAS: 2 traits were defined on hg17, 13 on hg18, and 54 on hg19. Panel **A** shows the distribution of number of associations per trait-tissue pair that can be computed; imputation dramatically increased the number of associations for hg17- and hg18-based traits. Some hg19-based traits exhibited a good number of computable associations after just a simple harmonization. Panel **B** shows, per trait-tissue pair, the distribution of median fraction of model SNPs present in the GWAS. It is nearly 1 for most trait-tissue pairs in the imputation scheme, ranging between 0.5 and 1 with the harmonization scheme. Panel **C** shows the number of colocalized, significantly associated genes that can be found after applying imputation and harmonization schemes. The gain of imputation for hg19 is less dramatic that in the other comparisons in this figure, given the conservative nature of the colocalization filter.

Fig. 5-A summarizes the increase in number of gene associations computed for every trait-tissue pair. For hg17- and hg18-based GWAS, the gain through summary-statistics imputation is almost threefold. Some hg19-based GWAS traits without imputation yield a good enough number of computable genes.

Fig. 5-B shows the distribution of median fraction of model SNPs also present in the GWAS, within each tissue-trait combination. Roughly 60% of models’ variants are present in hg17- and hg18-based GWAS without imputation; this percenteage is substantially higher for hg19-based GWAS without imputation. Imputing summary statistics increases this median percenteage to 100% on all tissue-trait combinations across the analyzed human genome release versions.

Fig. 5-C shows the increase in number of genes detected per trait. As in the previous panels, the increase is more noticeable for hg17- and hg18-based GWAS, while smaller for hg19-based studies.

Therefore, we recommend to always perform variant harmonization due to its low complexity and time requirements, followed by summary-statistics imputation if possible. For newer GWAS with modern sequencing and genotyping, summary-statistics imputation may not be as critical depending on their intersection with model variants.

## Discussion

Through extensive analysis of different model training schemes, we conclude that using fine-mapping information (from *dap-g*) and cross-tissue patterns (from *mashr*) improve transcriptome prediction models both for causal gene detection and prediction performance. These models (MASHR-M) yield more detections when integrating GWAS and eQTL studies, and show better prediction performance on an independent expression cohort. This method also exhibits superior performance when validating results in a silver standard of known gene-to-trait associations (OMIM database). We make all prediction models and results publicly available.

Special consideration must be paid to how well each model’s variants intersect GWAS’ variants. Fine-mapping-informed models are sparse and parsimonious. This could be a hurdle when the fine-mapped variants of import are missing or have low imputation quality in a GWAS, as is often the case with older studies. In this scenario we recommend harmonizing and imputing summary statistics to the models’ variants. The alternative is falling back to models such as CTIMP-M, defined on a robust set of variants available to most GWAS, at the cost of decreased performance (detection and prediction). EN-M additionally features some “built-in” redundacy: for a set of variants in LD among each other, they all tend to be included in a model with the effect spread between them.

While our recommended MASHR-M method offers several benefits compared to existing approaches, there is still room for improvement. Potential developments could rely on fine-mapping methods that jointly incorporate cross-tissue patterns, or consensus between different fine-mapping approaches. Also, epigenetic information has been shown to improve transcriptome prediction [33] {Zhang2019EpiXcan} as well. Future improvements should incorporate this epigenetic information and other biologically-informed annotations jointly.

Our validation in silver standards, especially our difficulty interpreting the results from the rare-variant-based silver standard, also illustrates the need for well-curated, large databases of known gene-to-phenotype associations to assess performance of either new or improved methods.

In conclussion, we present here a method for predicting the genetically regulated component of transcriptomic traits with superior performance both in terms of prediction performance and gene-trait association detection.

## Acknowledgements

We thank the donors and their families for their generous gifts of organ donation for transplantation, and tissue donations for the GTEx research project.

The consortium was funded by GTEx program grants: HHSN268201000029C (F.A., K.G.A., A.V.S., X.Li., E.T., S.G., A.G., S.A., K.H.H., D.Y.N., K.H., S.R.M., J.L.N.), 5U41HG009494 (F.A., K.G.A.), 10XS170 (Subcontract to Leidos Biomedical) (W.F.L., J.A.T., G.K., A.M., S.S., R.H., G.Wa., M.J., M.Wa., L.E.B., C.J., J.W., B.R., M.Hu., K.M., L.A.S., H.M.G., M.Mo., L.K.B.), 10XS171 (Subcontract to Leidos Biomedical) (B.A.F., M.T.M., E.K., B.M.G., K.D.R., J.B.), 10ST1035 (Subcontract to Leidos Biomedical) (S.D.J., D.C.R., D.R.V.), R01DA006227-17 (D.C.M., D.A.D.), Supplement to University of Miami grant DA006227. (D.C.M., D.A.D.), HHSN261200800001E (A.M.S., D.E.T., N.V.R., J.A.M., L.S., M.E.B., L.Q., T.K., D.B., K.R., A.U.), R01MH101814 (M.M-A., V.W., S.B.M., R.G., E.T.D., D.G-M., A.V.), U01HG007593 (S.B.M.), R01MH101822 (C.D.B.), U01HG007598 (M.O., B.E.S.), R01MH107666 (H.K.I.), P30DK020595 (H.K.I.). E.R.G. is supported by the National Human Genome Research Institute (NHGRI) under Award Number 1R35HG010718 and by the National Heart, Lung, and Blood Institute (NHLBI) under Award Number 1R01HL133559. E.R.G. has also significantly benefitted from a Fellowship at Clare Hall, University of Cambridge (UK) and is grateful to the President and Fellows of the college for a stimulating intellectual home. S.K.-H. is supported by the Marie-Sklodowska Curie fellowship H2020 Grant 706636. R.D.: R35GM124836, R01HL139865, 15CVGPSD27130014. D.M.J.: T32HL00782. Y.Pa. is supported by the NHGRI award R01HG10067. A.R.H. was supported by the Massachusetts Lions Eye Research Fund Grant. H.E.W. is supported by NHGRI R15HG009569. Computation was performed at the high performance cluster of the Center for Research Informatics at the University of Chicago, funded by the Biological Sciences Division and CTSA UL1TR000430. Additional Computation was performed with resources provided by the University of Chicago Research Computing Center.

We thank the International Genomics of Alzheimer’s Project (IGAP) for providing summary results data for these analyses. The investigators within IGAP contributed to the design and implementation of IGAP and/or provided data but did not participate in analysis or writing of this report. IGAP was made possible by the generous participation of the control subjects, the patients, and their families. The i–Select chips was funded by the French National Foundation on Alzheimer’s disease and related disorders. EADI was supported by the LABEX (laboratory of excellence program investment for the future) DISTALZ grant, Inserm, Institut Pasteur de Lille, Université de Lille 2 and the Lille University Hospital. GERAD was supported by the Medical Research Council (Grant n° 503480), Alzheimer’s Research UK (Grant n° 503176), the Wellcome Trust (Grant n° 082604/2/07/Z) and German Federal Ministry of Education and Research (BMBF): Competence Network Dementia (CND) grant n° 01GI0102, 01GI0711, 01GI0420. CHARGE was partly supported by the NIH/NIA grant R01 AG033193 and the NIA AG081220 and AGES contract N01–AG–12100, the NHLBI grant R01 HL105756, the Icelandic Heart Association, and the Erasmus Medical Center and Erasmus University. ADGC was supported by the NIH/NIA grants: U01 AG032984, U24 AG021886, U01 AG016976, and the Alzheimer’s Association grant ADGC–10–196728.

## Code and data availability

Genotype-Tissue Expression (GTEx) project’s raw whole transcriptome and genome sequencing data are available via dbGaP accession number phs000424.v8.p1. All processed GTEx data are available via GTEx portal. Imputed summary results, *enloc, coloc*, PrediXcan, MultiXcan, *dap-g*, prediction models, and reproducible analysis are available in https://github.com/hakyimlab/gtex-gwas-analysis and links therein.

## URLs

1000 Genomes Project Reference for LDSC, https://data.broadinstitute.org/alkesgroup/LDSCORE/1000G_Phase3_plinkfiles.tgz;

1000 Genomes Project Reference with regression weights for LDSC, https://data.broadinstitute.org/alkesgroup/LDSCORE/1000G_Phase3_weights_hm3_no_MHC.tgz;

BioVU, https://victr.vanderbilt.edu/pub/biovu/?sid=194;

eCAVIAR, https://github.com/fhormoz/caviar;

QTLEnrich, https://github.com/segrelabgenomics/eQTLEnrich;

flashr, https://gaow.github.io/mnm-gtex-v8/analysis/mashr_flashr_workflow.html#flash

Gencode, https://www.gencodegenes.org/releases/26.html;

GTEx GWAS subgroup repository, https://github.com/broadinstitute/gtex-v8;

GTEx portal, http://gtexportal.org;

Hail, https://github.com/hail-is/hail;

HapMap Reference for LDSC, https://data.broadinstitute.org/alkesgroup/LDSCORE/w_hm3.snplist.bz2;

LD score regression (LDSD regression), https://github.com/bulik/ldsc;

MetaXcan, https://github.com/hakyimlab/MetaXcan;

Mouse Phenotype Ontology, http://www.informatics.jax.org/vocab/mp_ontology;

NHGRI-EBI GWAS catalog, https://www.ebi.ac.uk/gwas/;

picard, http://picard.sourceforge.net/;

PLINK 1.90, https://www.cog-genomics.org/plink2;

PrediXcan, https://github.com/hakyimlab/MetaXcan;

pyliftover, https://pypi.org/project/pyliftover/;

Storey’s qvalue R package, https://github.com/StoreyLab/qvalue;

Summary GWAS imputation, https://github.com/hakyimlab/summary-gwas-imputation;

TORUS, https://github.com/xqwen/torus;

UK Biobank GWAS, http://www.nealelab.is/uk-biobank/;

UK Biobank, http://www.ukbiobank.ac.uk/;

## Methods

We executed all methods using open source software running in a high performance cluster. We release all of our code and the data analyzed in this paper to ease reproducibility and accessibility.

### GTEx data processing

We downloaded GTEx data for version 8 release from dbGAP (accession number phs000424.v8.p1). This data arises from 17382 RNA-seq samples from 54 tissues of 948 post-mortem subjects, aligned to the GRCh38 assembly. Primary and extended results generated by consortium members are available on the Google Cloud Platform storage accessible via the GTEx Portal (see URLs).

899 whole-genome sequencing (WGS) samples were analyzed, 68 of them at an average coverage of 30x on HiSeq200, and the rest on HiSeqX. 866 GTEx donors’ samples were included in the downstream variant call files (VCF), after excluding one each from 30 duplicate samples and 3 donors. Among these, 838 subjects with RNA-seq data were included for QTL mapping and analysis.

Whole transcriptome RNA-Seq data were aligned using STAR (v2.5.3.a; [34] {dobin:2013} For STAR index, GENCODE v26 was used with the sjdbOverhang 75 for 76-bp paired-end sequencing protocol. Default parameters were used for RSEM (see URLs; [35] {Li:2011}) index generation. GTEx utilized Picard (see URLs) to mark and remove potential PCR duplicates and RNA-SeQC [36] {DeLuca:2012dp} to process post-alignment quality control. RSEM was then used for per-sample transcript quantification. Subsequently, read counts were normalized between samples using TMM [37] {robinson:2010}. For eQTL analyses, latent factor covariates were calculated using PEER [38] {stegle:2010} as follows: 15 factors for N < 150 per tissue; 30 factors for 150 ≤N < 250; 45 factors for 250 ≤ N < 350; and 60 factors for N ≥ 350. Expression phenotypes were adjusted for unwanted variation using covariates such as gender, sequencing plaform and pcr protocol, the top 5 principal components from genotype data, and said PEER factors. Finally, fastQTL [39] {ongen:2016} was used for cis-eQTL mapping in each tissue. Only proteincoding, lincRNA, and antisense biotypes as defined by Gencode v26 were considered for further analyses. To study alternative splicing, GTEx applied LeafCutter (version 0.2.8; [20]{Li:2018cy}) using default parameters to quantify splicing QTLs in cis with intron excision ratios [2]{Aguet2019}.

We used the *dap-g [22]*{Wen2016}, *enloc [16]*{Wen2017} and *coloc [26]*{Giambartolomei2} results published in [2]{Aguet2019}.

### GTEx expression and splicing modelling

We used the same genotypes, phenotypes, covariates, gene annotations and variant annotations from the main GTEx analysis.

When building prediction models, we imposed an additional restriction: we used only samples of European ancestry for the sake of leveraging a well defined population LD structure. Only variants with MAF> 0.01 in these samples were included. We used 49 tissues with sample sizes ranging from 65 (Kidney Cortex) to 602 (Muscle Skeletal).

This ancestry restriction mitigated problems due to LD mismatch when integrating with most publicly available GWAS summary statistics, which are conducted on predominantly European populations. Prediction models in other ancestries are important, and we are currently dedicating substantial effort to creating and analyzing such models. However, non-European models are beyond the scope of this paper.

We only generated models for genes annotated in GENCODE v26 as protein coding, lncRNA or pseudogenes.

### Elastic Net models

We fitted an Elastic Net model for each gene-tissue pair with available adjusted expression data. We restricted the set of variants to those present in the HapMap 3 CEU track 40 {InternationalHapMap3Consortium2010} with MAF> 0.01. The motivation behind this choice was to restrict the analysis to a robust set of SNPs that has significant intersection with most publicly available GWAS summary statistics. For every gene, variants within 1Mb upstream of the gene’s transcription start site and 1Mb downstream of the transcription end site where used as explanatory variables for gene expression. We used the R package *glmnet [14]* {Friedman2010GLMNET}, with mixing parameter *α* = 0.5 and penalty parameter chosen through 10-fold cross validation.

Prediction performance was estimated using a nested cross-validation approach. Expression was predicted out-of-sample for each fold, with Elastic Net parameters estimated only within training data, and the correlations to observed values at each fold were combined via Fisher’s transformation and Stouffer’s method. Only those models with mean Pearson correlation across 10 folds *ρ* > 0.1 and nested cross-validated correlation test *p* < 0.05 were kept.

We refer to these models as EN-M.

### CTIMP models

We employed the CTIMP [11] {Hu2019} framework on the same data from EN-M models in the previous section. This method fits expression for a gene in multiple tissues simultaneously through a regularized linear model, using a Lasso penalty within each tissue and a group-Lasso penalty for cross-tissue patterns. As it internally uses genotypes from all samples available across all tissues, we expect improvements over EN-M to be larger for tissues of smaller sample size where EN-M deals with a less informative LD structure among variants.

We performed five-fold cross validation for model tuning and evaluation following the authors’ description. We computed cross-validated correlation measures across folds as in the previous method, and kept those models achieving the thresholds of cross-validated correlation *ρ* > 0.1 and p-value *p* < 0.05. As in EN-M, we restricted the model training to variants in the HapMap 3 CEU track with MAF> 0.01; this became necessary because using all variants proved too computationally expensive, since CTIMP consumes large amounts of memory and processing time. We briefly show in the Supplement (Supplementary Figures 4, 5, 6) that this additional restriction brings negligible effects in model training performance and prediction.

We refer to these models as CTIMP-M.

### Elastic Net informed by *dap-g* results

We also trained models via the Elastic Net algorithm using fine-mapping information to refine the list of variants to be used as explanatory variables, and lent more weight to variants with higher chances of affecting expression phenotypes. To this aim, we used *dap-g* ‘s posterior inclusion probability (PIP) of a variant affecting gene expression to select explanatory variables, without restricting to variants in the HapMap CEU track. For every gene, we used all variants in the gene’s cis-window with MAF> 0.01 and PIP> 0.01. Since *dap-g* groups variants in clusters according to LD, we kept the top variant (by PIP) per cluster to avert variable redundancy. Since we reasoned that more probable variants should bear more impact in the model’s outcome, we multiplied each variant’s penalty term in the Elastic Net regularization by a factor of 1 − PIP. We used the same thresholds from the previous subsections (*ρ* > 0.1 and p-value *p* < 0.05) to select models with acceptable prediction performance.

We refer to these models as DAPGW-M.

### *mashr*-based models

Finally we explored an entirely different algorithm to determine the prediction models. We executed multivariate adaptive shrinkage in R (*mashr*) [18] {Urbut2019} to estimate the models’ effect sizes by leveraging cross-tissue variations while allowing for sparse and possibly correlated effects in a Bayesian framework. We used *mashr* on the same set of variants from DAPGW-M models. We kept models only for eGenes and effect sizes only for variants with *PIP* > 0.01 (from *dap-g*) at each gene-tissue pair. Unfortunately, there is no natural prediction performance measure in this scenario as cross-validation was not performed.

We refer to these models as MASHR-M.

### GEUVADIS data processing

We used GEUVADIS LCL expression study for an independent validation of prediction performance. We obtained GEUVADIS expression data and sample information from the European Bioinformatics Institute web portal at https://www.ebi.ac.uk/. We obtained genotype data aligned to GRCh38 assembly from the International Genome Sample Resource web portal http://www.internationalgenome.org. We restricted data to individuals of European ancestry, yielding 341 samples.

For each one of the four previous model training schemes (EN-M, CTIMP-M, DAPGW-M, MASHR-M) we predicted expression through PrediXcan [8] {Gamazon2015} on GEU-VADIS genotypes using GTEX LCL models, and correlated predictions to observations.

### GWAS processing and integration

We examined 87 GWAS from a heterogeneous set of traits first presented in the GTEx v8 study [2, 23] {GTEx-GWAS-Companion, Aguet2019}. These traits were selected to support a phenome-wide study of the impact of gene regulation. Given the heterogeneous landscape of the GWAS, with intricate differences in data processing protocols and underlying human genome reference versions, it was necessary to make the GWAS variants homogeneous and compatible with those from the GTEx study.

First, the GWAS’ variants were harmonized to the GTEx study’s variants by mapping genomic coordinates via liftover [41] {haeussler:20019} (https://pypi.org/project/pyliftover) and keeping only variants with matching alleles. Then, GTEx variants with missing summary statistics for any GWAS were imputed with the BLUP method, a standard in the field [42] {Lee2013}.

We executed S-PrediXcan for each of 4 families of models (EN-M, CTIMP-M, DAPGW-M and MASHR-M) using 49 tissues, for a total of 17,052 (trait, model family, tissue) tuples. We integrated with *enloc* and *coloc* results published in [2] {Aguet2019}. When analying versatility of the models and GWAS preprocessing schemes, we used GWAS studies not belonging to the rapid GWAS study. This was decided because the rapid GWAS project has a common, homogeneous variant set that could dominate comparisons.

## Supporting information

### Restricting CTIMP variants to CEU HapMap track

When first attempting to run CTIMP package on GTEx data, we observed a significantly larger computational burden than the related Elastic Net method, up to two orders of magnitude larger in memory, and up to three in compute time for many genes. This scheme was too prohibitive and would had taken months to complete on the high performance cluster available to us.

To reduce the computational burden, we decided to restrict the explanatory variables to variants in the HapMap 3 CEU track [40] {InternationalHapMap3Consortium2010} with MAF> 0.01. This brought down the memory and processing consumption to only one order of magnitude larger than Elastic Net.

To verify that this technical restriction did not degrade performance too severely, we computed CTIMP models using all variants for chromosome 1, which took over 5 weeks to complete in our computation resource. We refer to these models as CTIMP-M-AS, which allowed comparison to CTIMP-M.

In Supplementary Figure 4-A we compare prediction performance *R*^2^ between CTIMP-M and CTIMP-M-AS. When both methods converge, they tend to agree. CTIMP-M-AS achieves better *R*^2^ on some genes. Supplementary Figure 4-B shows application of S-PrediXcan to one trait (ADIPOGEN) using only one tissue (Adipose Subcutaneous) for illustration purposes. Both methods perform similarly and this behavior is observed in all other tissue-trait combinations.

Supplementary Figure 5 compares CTIMP-M and CTIMP-M-AS models across the array of studied phenotypes. Supplementary Figure 5-A shows that both CTIMP-M and CTIMP-M-AS yield a similar number of models, with CTIMP-M-AS converging on additional genes as expected from its larger pool of explanatory variables. Supplementary Figure 5-B shows that the distribution of prediction performances is similar for both models.

Supplementary Figure 6 summarizes S-PrediXcan associations’ zscores in a similar fashion for a few sample traits, both using all associations (significant or not), and using only colocalized associations. We observe no significant gain in association strength using all variants (CTIMP-M-AS) compared to CEU HapMap variants (CTIMP-M).

### Intersection of GWAS and Model variants

EN-M and CTIMP-M are based on variants common in most publicly available GWAS. On the other hand, DAPGW-M and MASHR-M models’s intersection with a typical public GWAS is lower (see Supplementary Note). Therefore a tradeoff arises when integrating models with GWAS, posing the superior but more demanding MASHR-M models against the underperforming but robust EN-M or CTIMP-M models.

As an example of the importance of harmonizing and imputing GWAS summary statistics to the transcriptome reference data set (GTEx), consider two relatively new GWAS studies: blood traits from UK Biobank/INTERVAL and height from UK Biobank, without harmonizing nor imputing missing summary statistics. EN-M and CTIMP-M have over 97% of their variants present in said GWAS, while this percenteage drops to around 80% for DAPGW-M and MASHR-M. DAPGW-M shows a slightly higher intersection of variants than MASHR-M. Table 1 summarizes the number for variants present in each family, the subset among them present in the GWAS traits, and the fraction of these two numbers. In our experience, 80% is acceptable for running tools such as PrediXcan in some applications. However, when applying to older studies with potentially lower intersection between GWAS variants and the models’, imputing summary statistics for missing variants becomes necessary.

**Supplementary Table 1.** Intersection between prediction model and two GWAS studies. **family** is the model strategy, **gwas** is the study, **percent** measures the proportion of model snps present in the GWAS, **present** is the number of model snps present in the gwas, **n** is the total number of unique variants in the models.

| family | gwas | percent | present | n |
| --- | --- | --- | --- | --- |
| CTIMP-M | astle | 97% | 558457 | 575256 |
| CTIMP-M | ukb | 99% | 567246 | 575256 |
| EN-M | astle | 98% | 1445370 | 1476991 |
| EN-M | ukb | 99% | 1460231 | 1476991 |
| DAPGW-M | astle | 84% | 322950 | 386411 |
| DAPGW-M | ukb | 78% | 301318 | 386411 |
| MASHR-M | astle | 82% | 272282 | 331523 |
| MASHR-M | ukb | 76% | 251551 | 331523 |

To fully exploit the power available to the superior MASHR-M models, harmonizing becomes necessary in newer GWAS; imputation of missing sumamry statistics is also necessary in older GWAS.

## Supplementary Figures

**Supplementary Figure 1.**
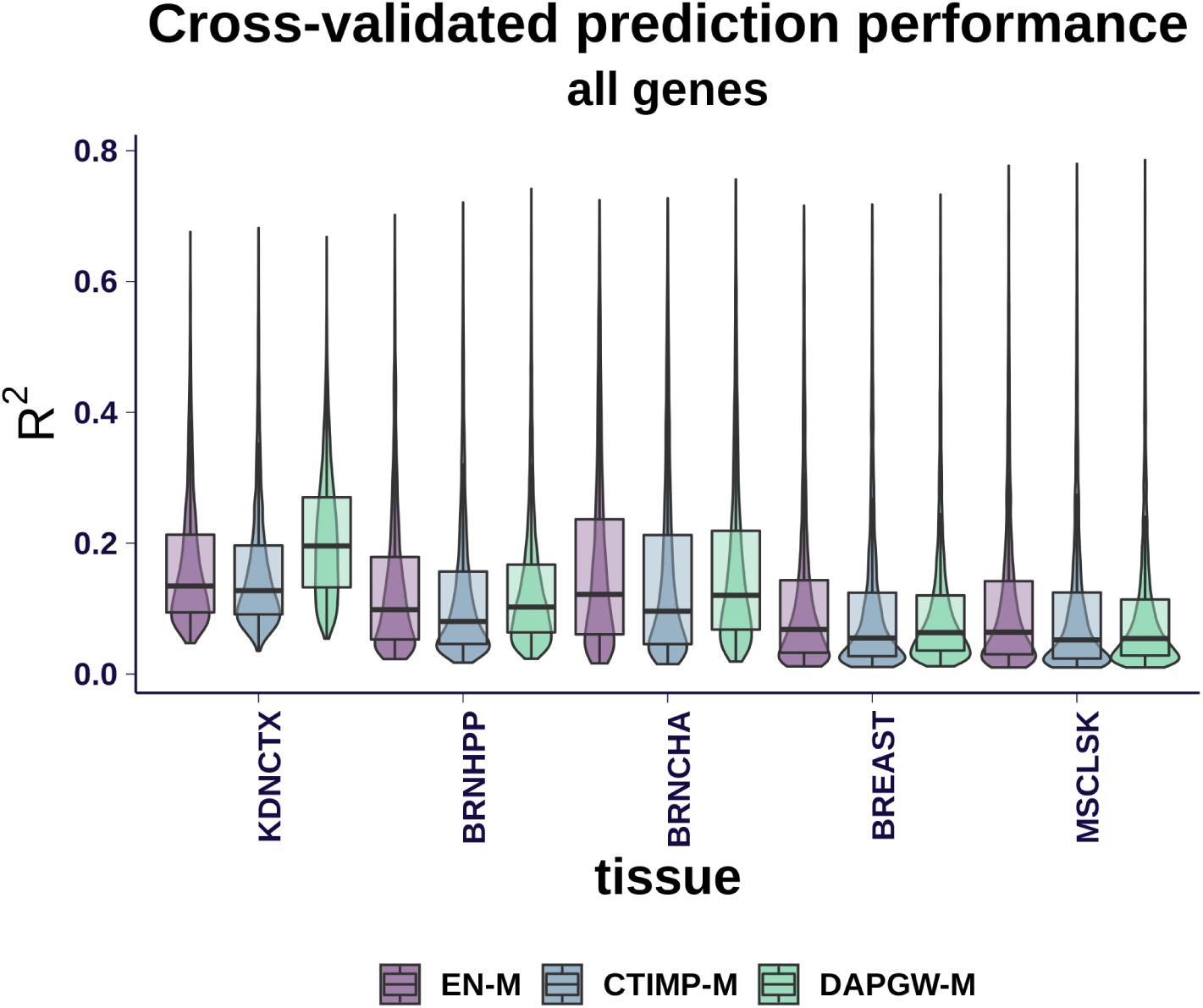
Cross-validated prediction performance is shown for all available models, on sample tissues ordered from smallest sample size to largest sample size. As sample size increased, we observed similar performances on the strategies shown. DAPGW-M is presented for illustration purposes; since it included an additional variable selection step using the same underlying data, it cannot be fairly compared to EN-M and CTIMP-M. We note that EN-M models produced the smallest number of models (Fig. 1-A), and 82% of them are in the intersection of models available to all strategies. These tend to be genes with higher heritability and thus easier to predict. In other words, EN-M models are generally available to CTIMP-M and DAPGW-M, and the intersection of all strategies is dominated by genes in EN-M. On the other hand, CTIMP-M and DAPGW-M yield viable models for additional genes that are harder to predict, where EN-M couldn’t converge to a proper model. In conclusion, CTIMP-M’s and DAPGW-M’s performance summary on all available genes was penalized by their convergence on genes with less signal or more complicated expression patterns. **Tissue abbreviations and sample size:** KDNCTX: Kidney - Cortex, n=65; BRNHPP: Brain - Hippocampus, n=150; BRNCHA: Brain - Cerebellum, n=188; BREAST: Breast - Mammary Tissue, n=337; MSCLSK: Muscle - Skeletal, n=602

**Supplementary Figure 2.**
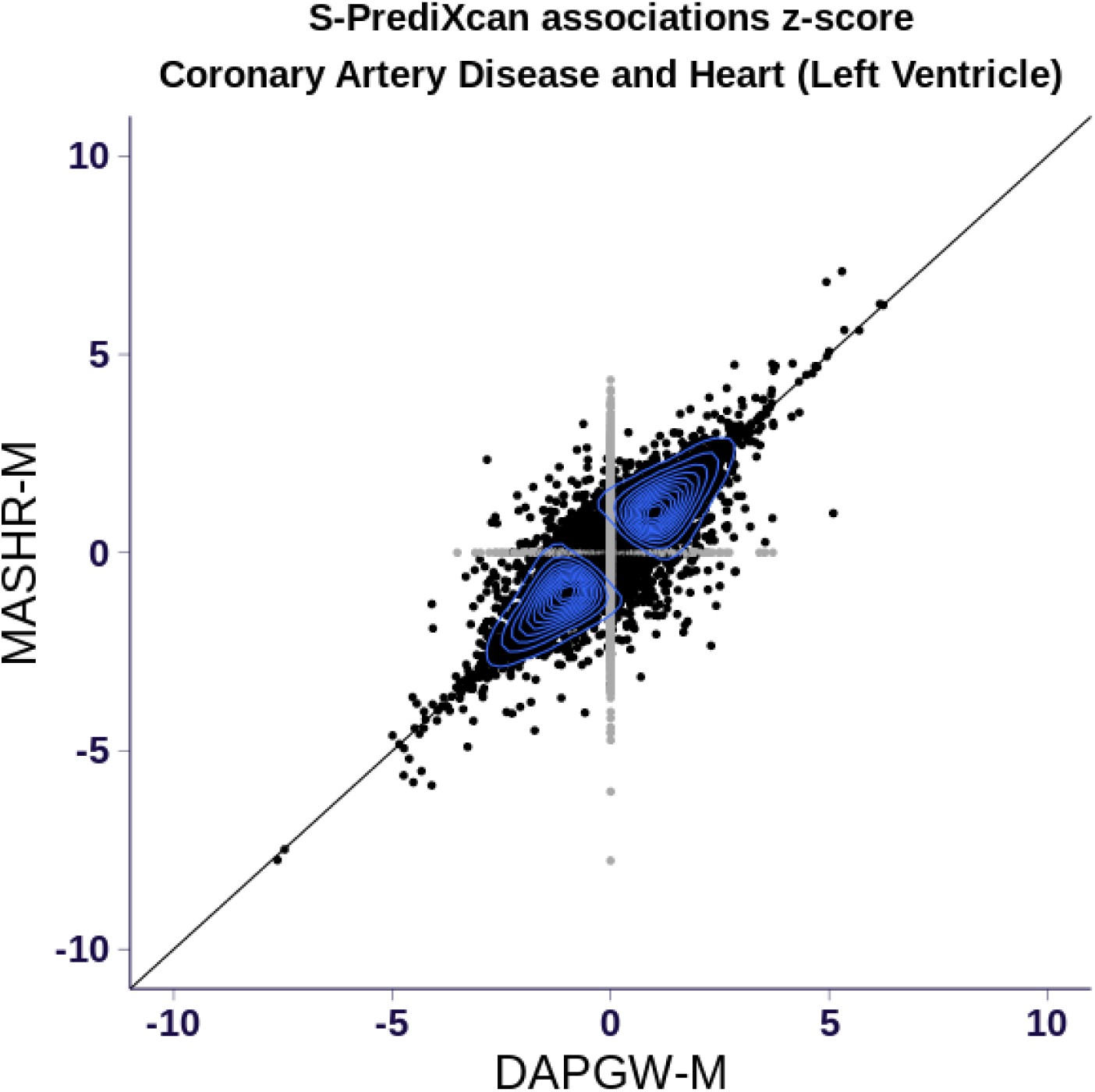
This figure compares S-PrediXcan associations for Coronary Artery Disease using Left Ventricle prediction models from DAPGW-M and MASHR-M. Black dots are associations present in both DAPGW-M and MASHR-M models, while gray dots are associations present in only one of them. For shared genes, both models tend to agree in association direction and magnitude. The concordance is higher for significant associations, while poorly associated genes might even disagree in direction. This behavior is common to all analyzed trait-tissue pairs.

**Supplementary Figure 3.**
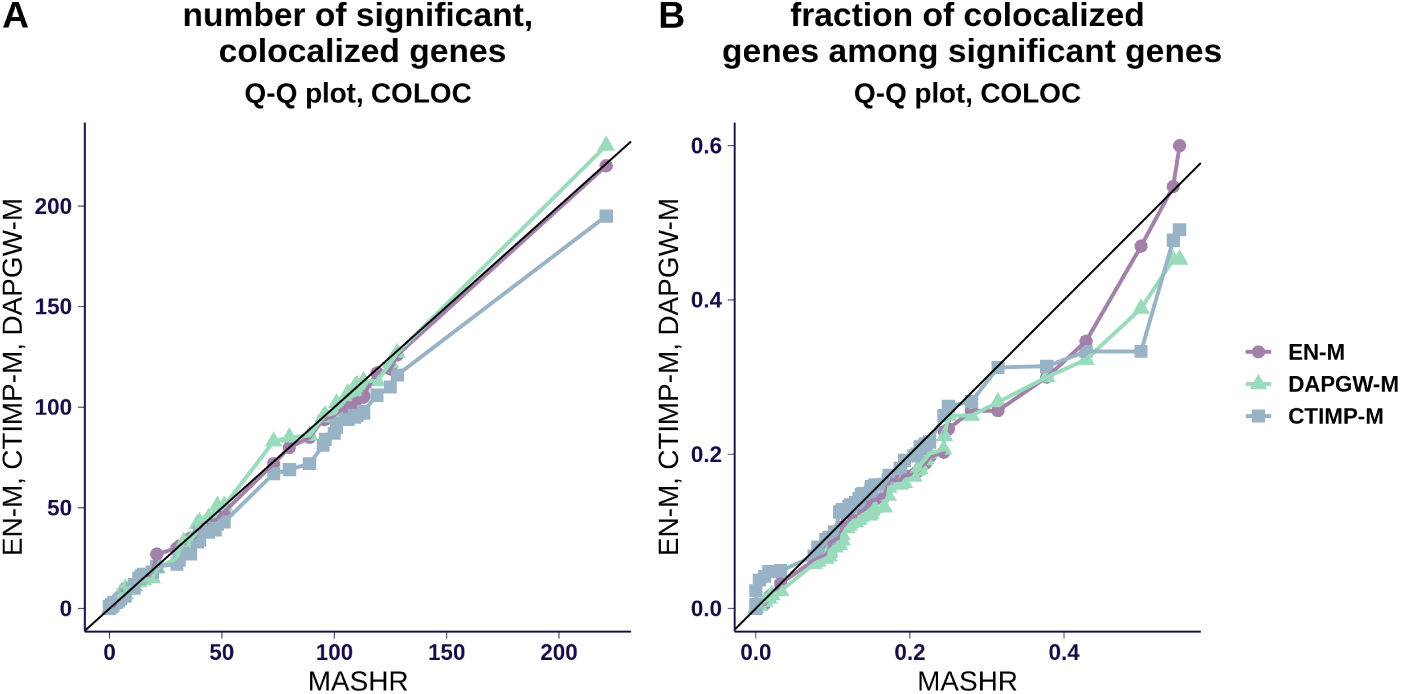
This figure compares the number of S-PrediXcan associations achieving *coloc*’s probability of colocalization PP4 > 0.5. In **panel A**, the number of detections based on *coloc* doesn’t clearly distinguish between the four methods. This is likely caused by *coloc*’s assumption of only one causal variant: since genes selected by *coloc* tend to have less allellic heterogeneity, they behave similarly across the four model strategies. In **panel B**, we observe that the fraction of significant genes that are also colocalized is slightly better for MASHR-M in general.

**Supplementary Figure 4.**
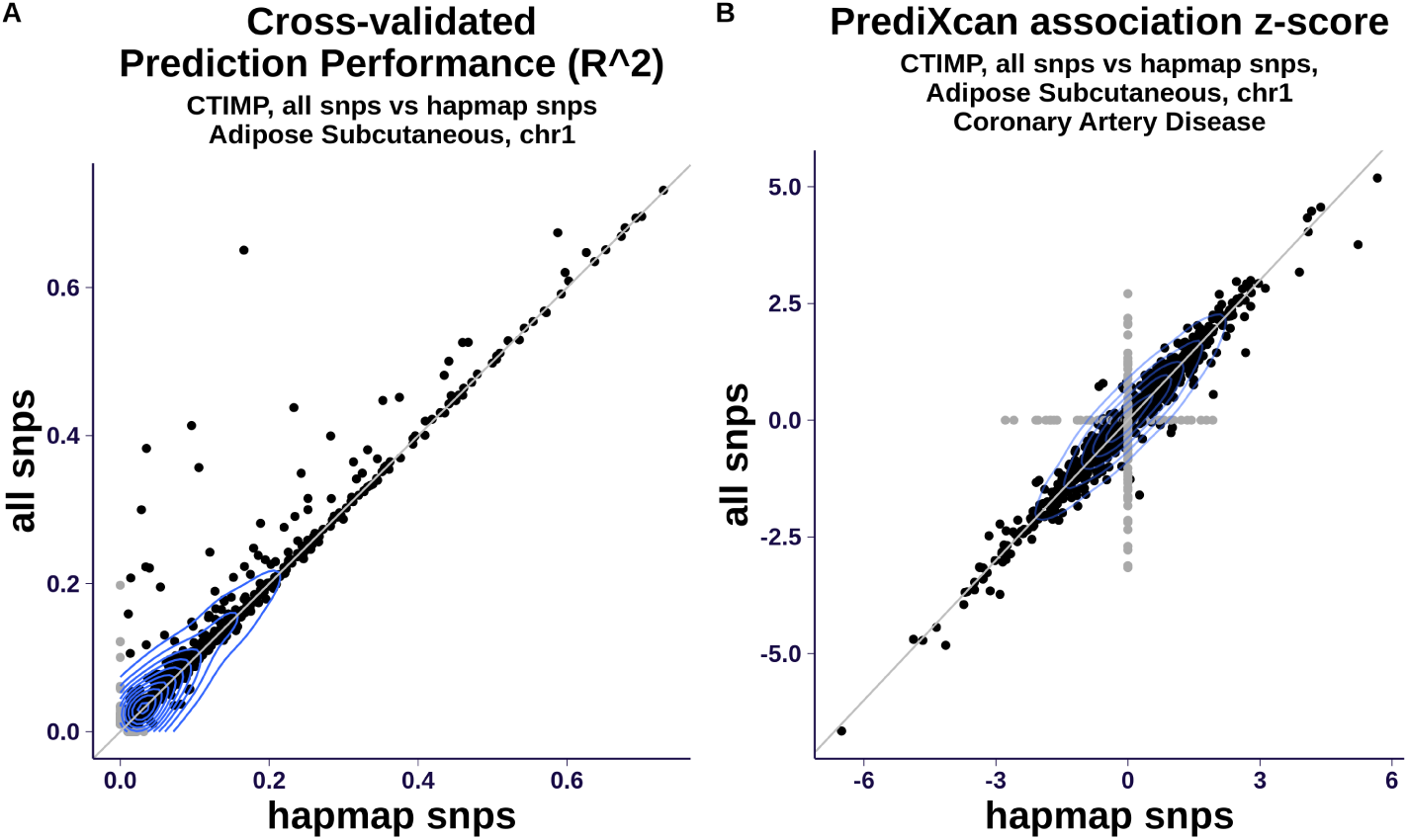
CTIMP Models using all variants (CTIMP-M-AS) vs HapMap variants (CTIMP-M). **Panel A** shows cross-validated prediction performance *R*^2^. When both methods converge, they tend to achieve similar prediction performances, with CTIMP-M-AS doing slightly better on some genes. **Panel B** compares S-PrediXcan associations for CTIMP-M and CTIMP-M-AS, showing a high level of agreement.

**Supplementary Figure 5.**
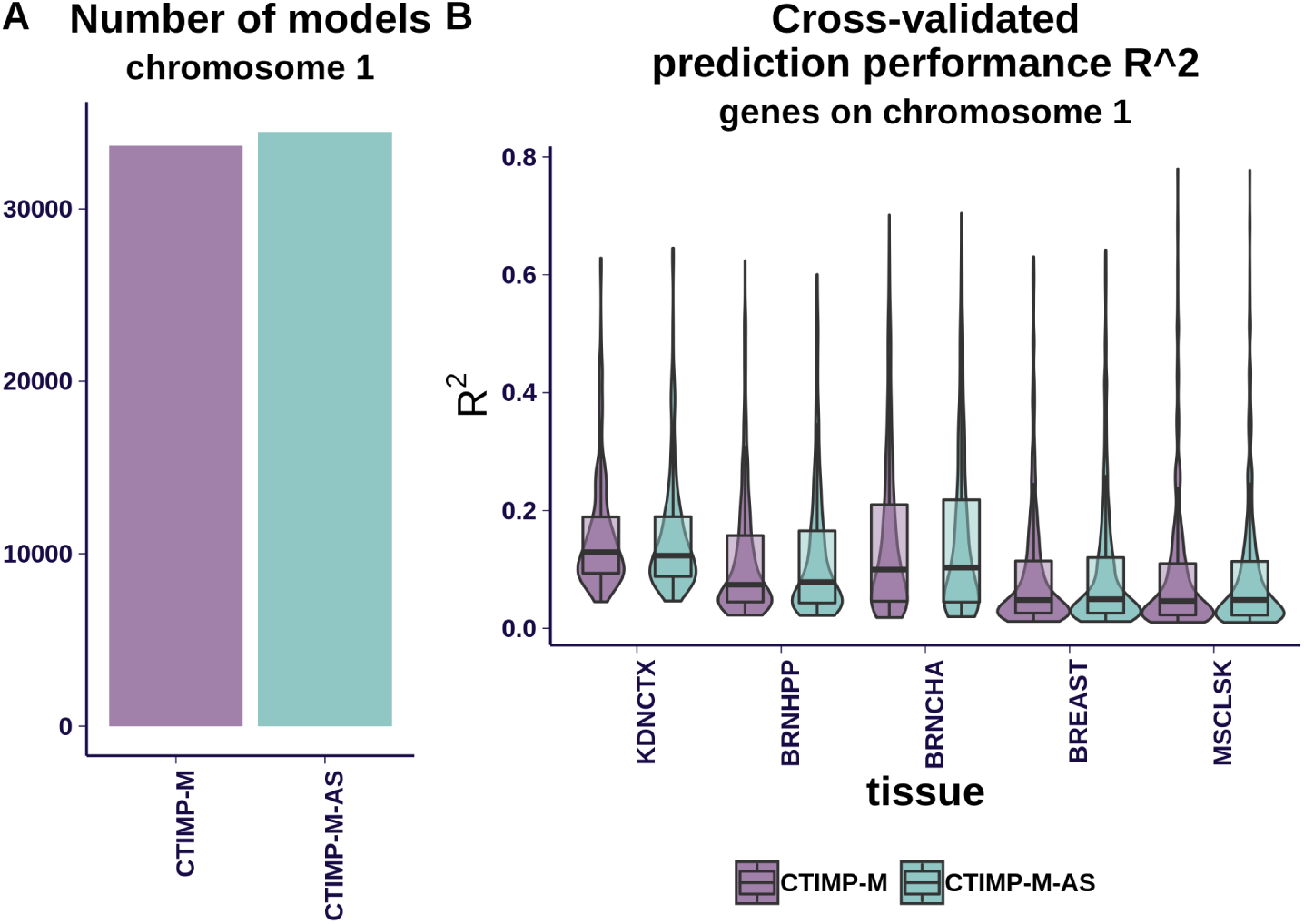
CTIMP Models summary. We compare here CTIMP using all variants (abbreviated as CTIMP-M-AS) to CTIMP using only HapMap variants (abbreviated as CTIMP-M), for genes in chromosome 1. We observed no significant difference between CTIMP-M and CTIMP-M-AS. **Panel A** shows the number of generated models for protein coding genes, pseudo genes and lncRNA. **Panel B** compares prediction performance for all gene-tissue pairs, for sample tissues (ordered from smallest sample size to largest sample size). The differences in performance are negligible.

**Supplementary Figure 6.**
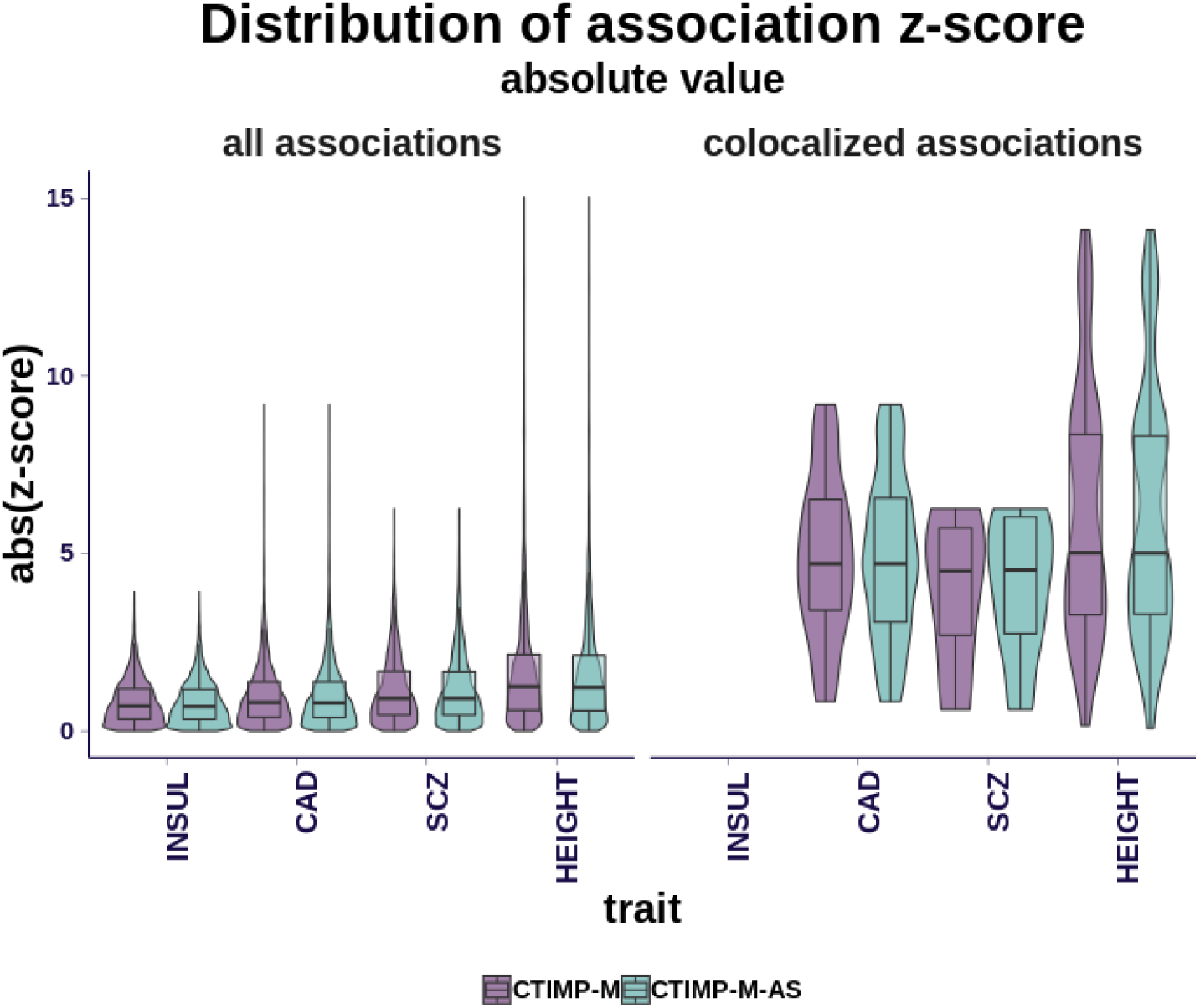
CTIMP’s PrediXcan associations for 4 sample traits from CTIMP models using all variants (abbreviated as CTIMP-M-AS) to CTIMP using only HapMap variants (abbreviated as CTIMP-M), for genes in chromosome 1. Both panels show absolute values of association z-score for gene-tissue pairs, to Fasting Insulin (INSUL), Coronary Artery Disease (CAD), Schizophrenia (SCZ) and Height (HEIGHT); traits are ordered from lowest to highest number of uniquely associated genes. The left panel shows all associations, whereas the right panel shows colocalized associations. We observed no significant difference between CTIMP-M and CTIMP-M-AS.

